# Plasmid transfer and loss probabilities interactively determine antibiotic resistance spread in surface-associated bacterial biomass

**DOI:** 10.64898/2026.01.08.698332

**Authors:** Josep Ramoneda, Deepthi P. Vinod, Yinyin Ma, Chujin Ruan, Julian Schmidt, Michael Manhart, Daniel C. Angst, David R. Johnson

## Abstract

Plasmid transfer among bacteria is an evolutionary driver of the spread of antibiotic resistance. Surface-associated bacterial biomass is a hotspot for plasmid transfer due to the dense spatial packing of cells. Compared to plasmid dynamics within a single biomass unit, the determinants of plasmid transfer between discrete units are less understood. Yet, discrete units routinely physically collide with each other as they grow, such as in sparse biofilms. Here, we used individual-based modelling to quantitatively predict how the probabilities of plasmid transfer and loss affect plasmid spread between discrete bacterial colonies as they grow and physically collide into each other. We found that these factors have interactive effects on the extent of plasmid transfer and the subsequent proliferation of transconjugants. We further found that these effects are controlled by the spatial distance between the colliding colonies and the spatial positioning of plasmid-carrying cells along the collision boundary. We then experimentally tested our modelling predictions using strains of *Pseudomonas stutzeri* and *Escherichia coli* that can exchange an antibiotic resistance-encoding plasmid. Our study reveals that plasmid transfer between colliding biomass units is determined by the complex interplay between plasmid transfer, plasmid loss, and spatial constraints, expanding our understanding of plasmid dynamics.

## Introduction

Surface-associated bacterial biomass such as biofilms, bio-aggregates, and colonies are pervasive on our planet [1], drive all major biogeochemical cycles [2, 3], and affect human health and disease [4–6]. They are engineered for diverse biotechnological applications such as water decontamination and biofuel production [7, 8], but also cause persistent infections in animal tissues and contaminate medical devices [9]. Within surface-associated biomass, the dense spatial packing and sessile state of bacterial cells can promote the contact-dependent horizontal transfer of plasmids and their associated genes [10]. This has serious consequences for the emerging antibiotic resistance crisis, as antibiotic resistance-encoding plasmids can cause persistent resistant bacterial populations in human and environmental microbiomes, posing a threat to global health [11, 12]. While critically important, our understanding of how the spatial features of (and spatial constraints acting on) surface-associated biomass direct the spread of antibiotic resistance-encoding plasmids remains at its infancy.

Despite the recognition that surface-associated bacterial biomass is a hotspot for plasmid transfer, the majority of plasmid transfer is postulated to occur along the outer edges of the biomass [13]. This is because plasmids generally only invade into and subsequently proliferate within metabolically active cells, which are usually those cells located at the outer edges of the biomass where unoccupied space and nutrients that enter from the periphery are plentiful [14]. This expectation is supported by empirical observations in systems such as the mouse gut [15], where plasmid transfer typically occurs only along the periphery of the mucus layer covering epithelial cells. Recent studies with bacterial colonies growing across nutrient-rich agar surfaces, however, demonstrate that plasmid transfer can be pervasive within the entire biomass [16, 17]. To better understand plasmid dynamics within surface-associated biomass, it is necessary to i) delineate plasmid transfer occurring within the biomass from transfer occurring along its periphery, and ii) determine whether plasmid transfer at these locations is driven by the same or different mechanisms.

Plasmid dynamics within a single patch of surface-associated biomass are determined by how easily plasmids transfer between and persist among the component populations, which is primarily controlled by the likelihood that cells acquire plasmids from each other when in close spatial proximity (i.e. the plasmid transfer probability), the likelihood that plasmids are lost during cell division (i.e. the plasmid loss probability), and the strength of selection on plasmid-encoded traits [18, 19]. Thus, plasmid loss can be offset by high plasmid transfer probabilities, leading to the persistence of antibiotic resistance even in the absence of positive selection [20]. Since plasmid transfer and loss probabilities determine the load and distribution of plasmids over time, we expect these mechanisms to be important determinants of plasmid spread within and between surface-associated biomass units.

In addition to nutrient availability, the spatial arrangement of cells within a unit of surface-associated biomass is another important determinant of plasmid transfer and spread [14, 16, 17]. During surface-associated biomass growth, the component populations typically spatially segregate from each other as a consequence of stochastic drift along the periphery of the biomass [21]. This process has important effects on diversity [22, 23], stability [24] and functioning [25]. Plasmids transfer more extensively when the component populations are highly spatially intermixed, which can be ascribed to the greater number of cell-cell contacts between plasmid-free and -carrying cells that are necessary for plasmid transfer [13, 26, 27]. Frequent disturbance can also promote plasmid transfer by causing the spatial reorganization of cells and creating new cell-cell contacts between otherwise spatially-segregated populations [14, 28].

While the processes governing plasmid transfer within a single unit of surface-associated biomass are now becoming clear, surfaces are generally not colonized by a single contiguous biomass unit but rather by many spatially discrete biomass units (e.g., sparse biofilms or dispersed colonies), where each biomass unit lies adjacent to others and dynamically expands and contracts in size as a consequence of growth and death [29, 30]. Despite the pervasiveness of such sparsely colonized surfaces, there is little information on the processes affecting plasmid transfer between spatially discrete biomass units [13, 31, 32]. Scenarios where neighboring biomass units grow and eventually collide into each other are likely common in systems such as on medical implants [33, 34], dental plaque [35, 36], and wound infections [37]. The processes of growth, collision and retraction are more prominent when the system is periodically exposed to disturbances such as antibiotics, where antibiotic administration can drastically reduce the population sizes of sensitive individuals. This can increase biomass sparseness while also exacerbating the subsequent spread of plasmid-encoded antibiotic resistance during growth and recovery by imposing a positive selection pressure [38, 39]. Understanding the mechanisms governing the transfer of antibiotic resistance-encoding plasmids during physical collisions of discrete biomass units after such disturbances, in particular the role of plasmid transfer and loss probabilities in relation to spatial features, would help address the emerging antibiotic resistance crisis by improving our ability to predict plasmid dynamics and fate.

In this study, we investigated the determinants of the horizontal transfer of an antibiotic resistance-encoding plasmid during the growth and physical collision of discrete biomass units. To accomplish this, we adapted an individual-based model to simulate the growth and physical collision of discrete biomass units. We then used the model to establish quantitative predictions about how plasmid transfer and loss probabilities affect the spread of a conjugative plasmid between discrete biomass units located at different initial distances from each other using a full-factorial design (1500 simulations). We additionally used the model to investigate the influence of spatial features of the collision boundary between discrete biomass units on plasmid spread. Finally, we designed an experimental system in which a plasmid donor colony, consisting of a pair of fluorescently labelled strains of the bacterium *Pseudomonas stutzeri* A1601 that carry the self-transmissible conjugative plasmid pAR145, grows and eventually collides into an *Escherichia coli* potential recipient colony. pAR145 encodes chloramphenicol resistance and cyan fluorescent protein, thus enabling us to track and quantify it across space and time. By physically colliding the two-strain *P. stutzeri* plasmid donor colony with the one-strain *E. coli* potential recipient colony, we could test the model predictions by quantifying the extent of plasmid transfer as a function of the initial distance between the growing colonies.

## Results

### Individual-based modelling framework

We used the individual-based biophysical modelling framework of CellModeller [40] to test the influence of plasmid transfer and loss probabilities on plasmid spread during the growth and collision of discrete bacterial colonies in a full factorial manner. We simulated two colonies composed of distinct cell-type backgrounds: a plasmid donor colony composed of a 1:1 mixture of plasmid-carrying and -free cells and a potential recipient colony composed solely of plasmid-free cells. For our simulations, we set the plasmid-carrying cells of the plasmid donor colony to have a 17% lower growth rate than the plasmid-free cells (referred to as the plasmid cost). This plasmid cost is based on our experimental measurements of the reduction in the growth rate caused by carrying the pAR145 plasmid (Supplementary Table 1; Supplementary Figs. 1,2). We applied this plasmid cost to all transconjugant cells emerging in the plasmid donor colony. We did not apply a plasmid cost to transconjugant cells emerging in the potential recipient colony, which is again based on our experimental measurements (Supplementary Figure 3). Using this modelling approach, we could successfully quantify plasmid dynamics in the plasmid donor colony and capture the formation of transconjugant cells within the potential recipient colony upon colony collision (Fig. 1A,F).

**Figure 1.**
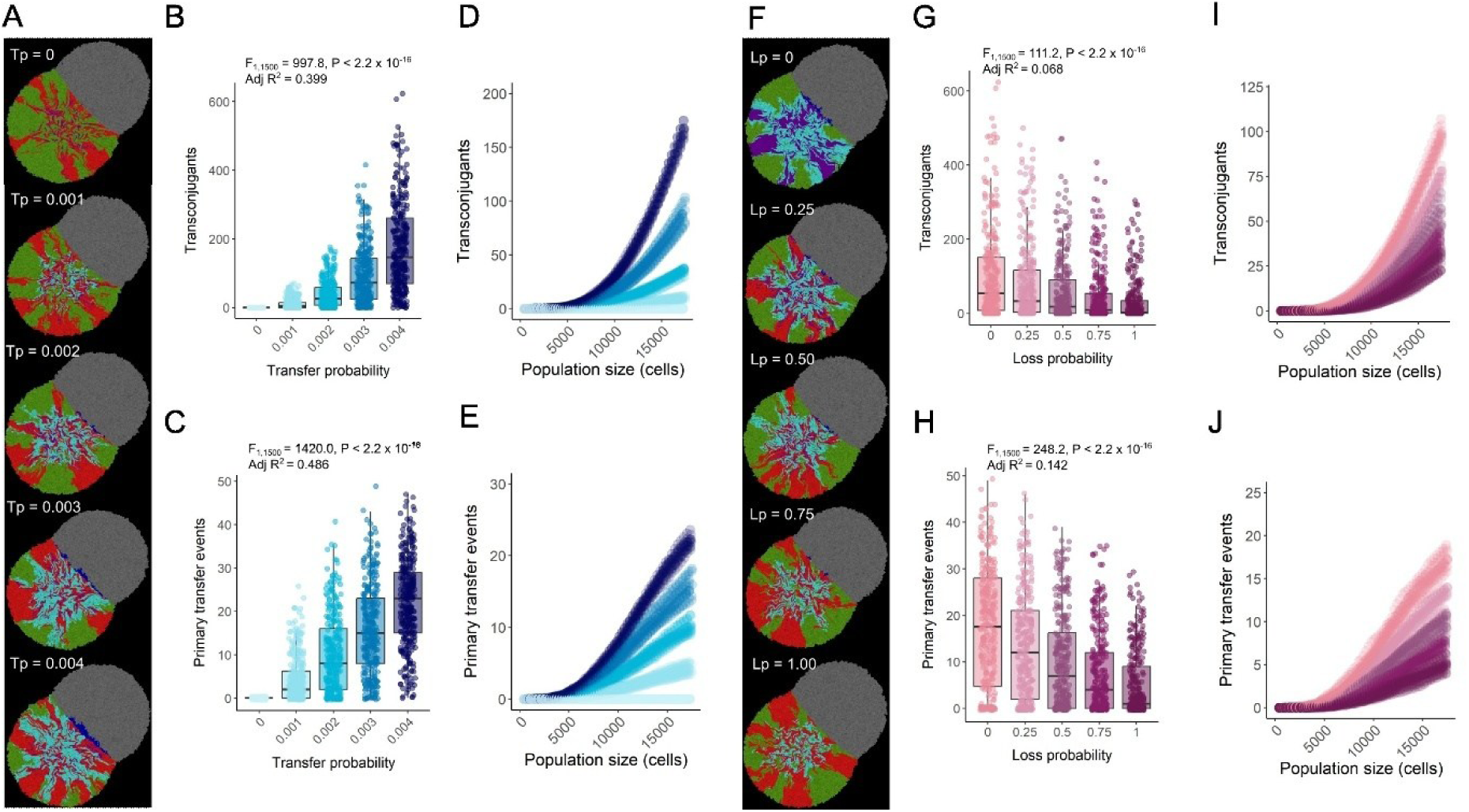
Effect of plasmid transfer and loss probabilities on plasmid spread between colliding colonies. (A and F) Representative individual-based simulations where the initial inoculum for the plasmid donor colony consisted of a 1:1 mixture of plasmid-carrying and -free cells and the inoculum for the potential recipient colony consisted of a single population of plasmid-free cells (grey) that can receive the plasmid upon collision (blue). T_p_ and L_p_ values at the top left of the simulation images are the plasmid transfer and loss probabilities for the simulations in (A) and (F), respectively. (B) Total number of transconjugant cells (blue) at the end of the simulations as a function of the plasmid transfer probability. (C) Total number of primary plasmid transfer events between the plasmid donor and potential recipient colonies at the end of the simulations as a function of the plasmid transfer probability. (D) Total number of transconjugant cells (blue) as a function of the total population size, colored by the plasmid transfer probability as defined in (B). (E). Total number of primary plasmid transfer events between the plasmid donor and potential recipient colonies as a function of the total population size, colored by the plasmid transfer probability as defined in (B). (G) Total number of transconjugant cells (blue) at the end of the simulations as a function of the plasmid loss probability. H. Total number of primary plasmid transfer events between the plasmid donor and potential recipient colonies at the end of the simulations as a function of the plasmid loss probability. I. Total number of transconjugant cells (blue) as a function of the total population size, colored by the plasmid loss probability as defined in (G). (J) Total number of primary plasmid transfer events between the plasmid donor and potential recipient colonies as a function of the total population size, colored by the plasmid loss probability as defined in (G). For (A), there is a probability of 0.5 at each cell division that plasmid-carrying cells will lose the plasmid. For (F), there is a probability of 0.003 that plasmid-carrying cells will transfer the plasmid to adjacent plasmid-free cells upon contact. Statistics in (B), (C), (G), and (H) are one-way ANOVA tests with the (B, C) plasmid transfer probability or (G, H) plasmid loss probability as the sole explanatory variable (n = 10).

### Contributions of plasmid transfer and loss to plasmid spread upon colony collision

We used the total number of transconjugant cells in the potential recipient colony and the total number of primary plasmid transfer events between cells within the plasmid donor and potential recipient colonies as descriptors of plasmid spread. We also quantified secondary plasmid transfer events (i.e. plasmid transfer between transconjugants and plasmid-free cells in the potential recipient colony). We quantified these descriptors at the same final population size across all simulations. We found that both the plasmid transfer and loss probabilities affect these quantities, with a predominant influence of the plasmid transfer probability (Fig. 1A,F). Specifically, the total number of transconjugant cells was strongly positively affected by the plasmid transfer probability (one-way ANOVA test; *F_1, 1500_* = 997.8, *R^2^_adj_* = 0.399, *P* < 2.2 x 10^-16^, n = 10) (Fig. 1B) and weakly negatively affected by the plasmid loss probability (one-way ANOVA test; *F_1, 1500_* = 111.2, *R^2^_adj_* = 0.068, *P* < 2.2 x 10^-16^, n = 10) (Fig. 1G). Likewise, the numbers of primary plasmid transfer events between colonies were significantly positively affected by the plasmid transfer probability (one-way ANOVA test; *F_1, 1500_* = 1420.0, *R^2^_adj_* = 0.486, *P* < 2.2 x 10^-16^, n = 10) (Fig. 1C) and negatively affected by the plasmid loss probability (one-way ANOVA test; *F_1, 1500_* = 248.2, *R^2^_adj_* = 0.142, *P* < 2.2 x 10^-16^, n = 10) (Fig. 1H). We verified that these effects remain valid when analyzing the data at a single common simulation time step of 640 rather than a single common population size (Supplementary Fig. 4). We also found that the effects of the plasmid transfer and loss probabilities on the total number of secondary plasmid transfer events were similar to the effects on the total number of transconjugant cells. Secondary plasmid transfer events within the potential recipient colony were significantly positively affected by the plasmid transfer probability (one-way ANOVA test; *F_1, 1500_* = 937.9, *R^2^_adj_* = 0.385, *P* < 2.2 x 10^-16^, n = 10) (Supplementary Fig. 5A) and weakly negatively affected by the plasmid loss probability (one-way ANOVA test; *F_1, 1500_* = 85.67, *R^2^_adj_* = 0.053, *P* < 2.2 x 10^-16^, n = 10; Supplementary Fig. 5C).

We next tracked the formation of new transconjugant cells during colony growth over time. The total number of transconjugant cells exhibited an exponential increase throughout the simulations across the tested plasmid transfer and loss probabilities (Fig. 1D,I). When we evaluated the number of primary plasmid transfer events along the collision boundary, we found that it increased linearly from the beginning of colony growth but had a decelerating trend beginning at a colony size of ca. 12,000 cells independent of the plasmid transfer and loss probabilities (Fig. 1E,J). Over time, the accumulation of secondary plasmid transfer events had qualitatively similar exponentially increasing trends as the accumulation of new transconjugants (Supplementary Fig. 5B,D).

### Effect of initial distance between colliding colonies on plasmid spread

Using our modelling framework, we quantified the effect of the initial distance between the plasmid donor and potential recipient colonies on plasmid spread. We found that the initial distance had a strong effect on plasmid spread, where shorter distances increased plasmid spread (Fig. 2A). The initial distance had a stronger negative effect on the total number of transconjugant cells (one-way ANOVA test; *F_1, 1500_* = 51.98, *R^2^_adj_* = 0.145, *P* < 2.2 x 10^-16^, n = 10) (Fig. 2B) than on the total number of primary plasmid transfer events between colonies (one-way ANOVA test; *F_1, 1500_* = 31.28, *R^2^_adj_* = 0.092, *P* < 2.2 x 10^-16^, n = 10) (Fig. 2C). Again, we verified that these effects remain valid when analyzing the data at a single common simulation time step of 640 rather than a single common population size (Supplementary Fig. 6). The effect of initial distance on the total number of secondary plasmid transfer events within the potential recipient colony was again comparable to the effects on the total number of transconjugant cells, where longer initial distances had a significant negative effect on secondary plasmid transfer (one-way ANOVA test; *F_1, 1500_* = 60.93, *R^2^_adj_* = 0.167, *P* < 2.2 x 10^-16^, n = 10) (Supplementary Fig. 7A).

**Figure 2.**
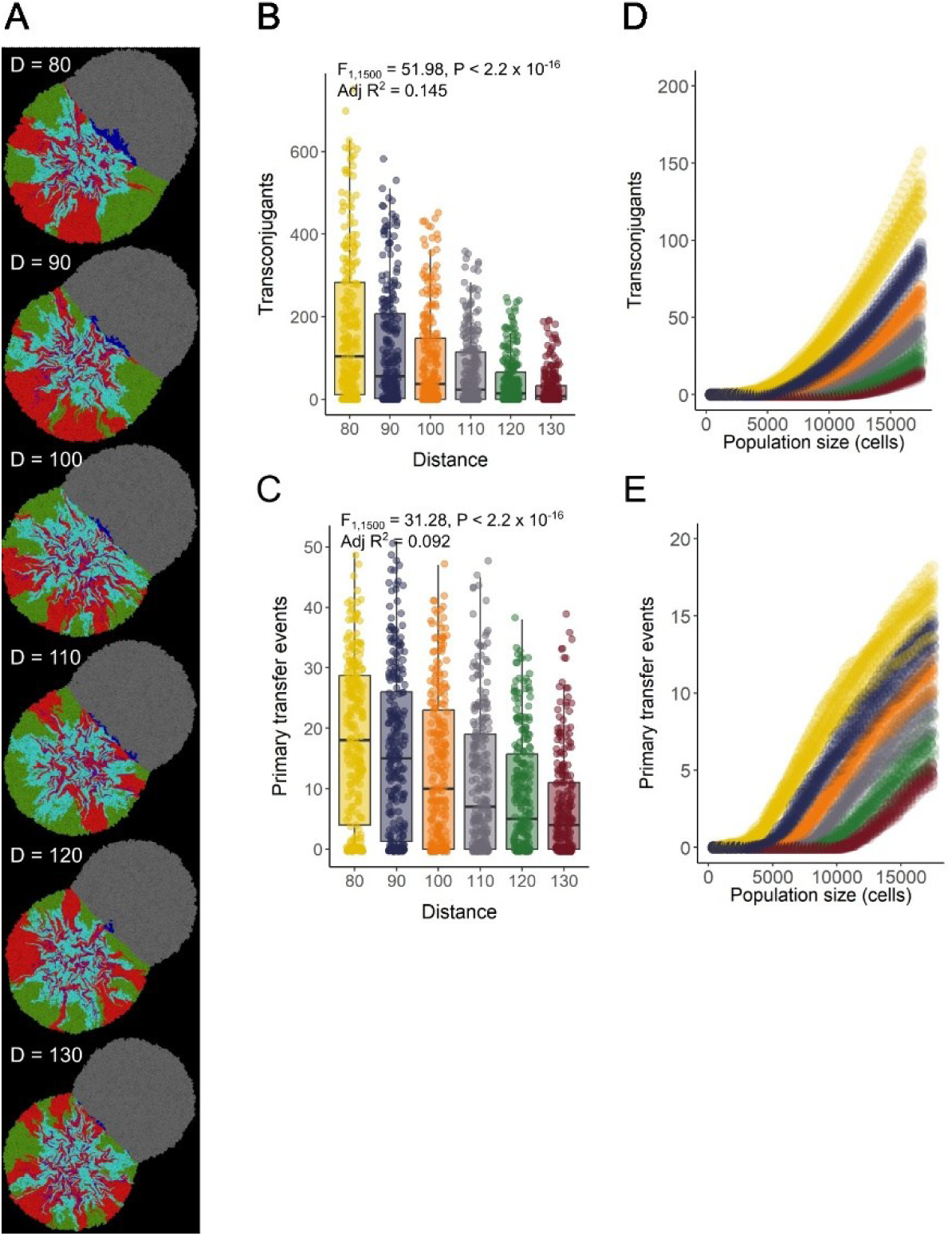
Effect of the initial distance between colliding colonies on plasmid spread. (A) Representative individual-based simulations where the initial inoculum for the plasmid donor colony consisted of a 1:1 mixture of plasmid-carrying and -free cells and the inoculum for the potential recipient colony consisted of a single population of plasmid-free cells (grey) that can receive the plasmid upon collision (blue). D_i_ values at the top left of the simulation images are the initial distances between the centroids of the inocula. (B) Total number of transconjugant cells (blue) at the end of the simulations as a function of the initial distance between inocula. (C) Total number of primary plasmid transfer events between the plasmid donor and potential recipient colonies at the end of the simulations as a function of the initial distance between inocula. (D) Total number of transconjugant cells (blue) as a function of the total population size. (E) Total number of primary plasmid transfer events between the plasmid donor and potential recipient colonies as a function of the total population size. For (A), there is a probability of 0.5 at each cell division that a plasmid-carrying cell will lose its plasmid and a probability of 0.003 that a plasmid-carrying cell will transfer the plasmid to an adjacent plasmid-free cell upon contact. Statistics in (B) and (C) are one-way ANOVA tests with initial distance as the sole explanatory variable (n = 10).

The initial distance had no effect on the shape of the curve describing the accumulation of new transconjugant cells or primary plasmid transfer events over time. The total number of new transconjugant cells again displayed a continuous and non-decelerating increase throughout the simulations regardless of the initial distance between colonies (Fig. 2D), while the total number of primary plasmid transfer events had a slightly decelerating trend beginning at a colony size of ca. 12,000 cells (Fig. 2E). The temporal trend of accumulation of secondary plasmid transfer events in the potential recipient colony was again exponential as we observed for the accumulation of transconjugant cells (Supplementary Fig. 7B).

### Initial distance determines the relative importance of plasmid transfer and loss in plasmid spread

Given the strong effect of the initial distance between colonies on plasmid spread between them, we quantitatively described how this initial distance determines the relative importance of plasmid transfer and loss on plasmid spread. We postulated that longer initial distances require longer times for colonies to collide, which provides more time to accumulate plasmid transfer and loss events within the plasmid donor colony. We found that the plasmid transfer probability is the main determinant of both the total number of transconjugant cells and the total number of primary plasmid transfer events between colonies regardless of the initial distance (Fig. 3A,B). However, the initial distance had a significant effect on the relative importance of these two processes for determining the total number of transconjugant cells (full model *R^2^_adj_* = 0.840, *P* < 2.2 x 10^-16,^ N = 1500) (Supplementary Table 2; Supplementary Table 3), the number of primary plasmid transfer events (full model *R^2^_adj_* = 0.816, *P* < 2.2 x 10^-16^, N = 1500) (Supplementary Table 2; Supplementary Table 3), and the number of secondary plasmid transfer events (full model *R^2^_adj_* = 0.819, *P* < 2.2 x 10^-16^, N = 1500) (Supplementary Table 4). We found that longer initial distances increased the relative importance of the plasmid loss probability over the plasmid transfer probability on plasmid spread (Fig. 3C). For the total number of transconjugant cells, the relative importance of the plasmid loss probability increased from 1.8% at an initial distance of 80 (two-way ANOVA test; F*_1,250_* = 118.53, *P* < 2.2 x 10^-16^, n = 10) to 5.2% at an initial distance of 130 (two-way ANOVA test; F*_1,250_* = 125.03, *P* < 2.2 x 10^-16^, n = 10). Correspondingly, the relative importance of the plasmid transfer probability decreased from 70.7% at an initial distance of 80 (two-way ANOVA test; F*_1,250_* = 1368.85, *P* < 2.2 x 10^-16^, n = 10) to 55.1% at an initial distance of 130 (two-way ANOVA test; F*_1,250_* = 286.33, *P* < 2.2 x 10^-16^, n = 10).

**Figure 3.**
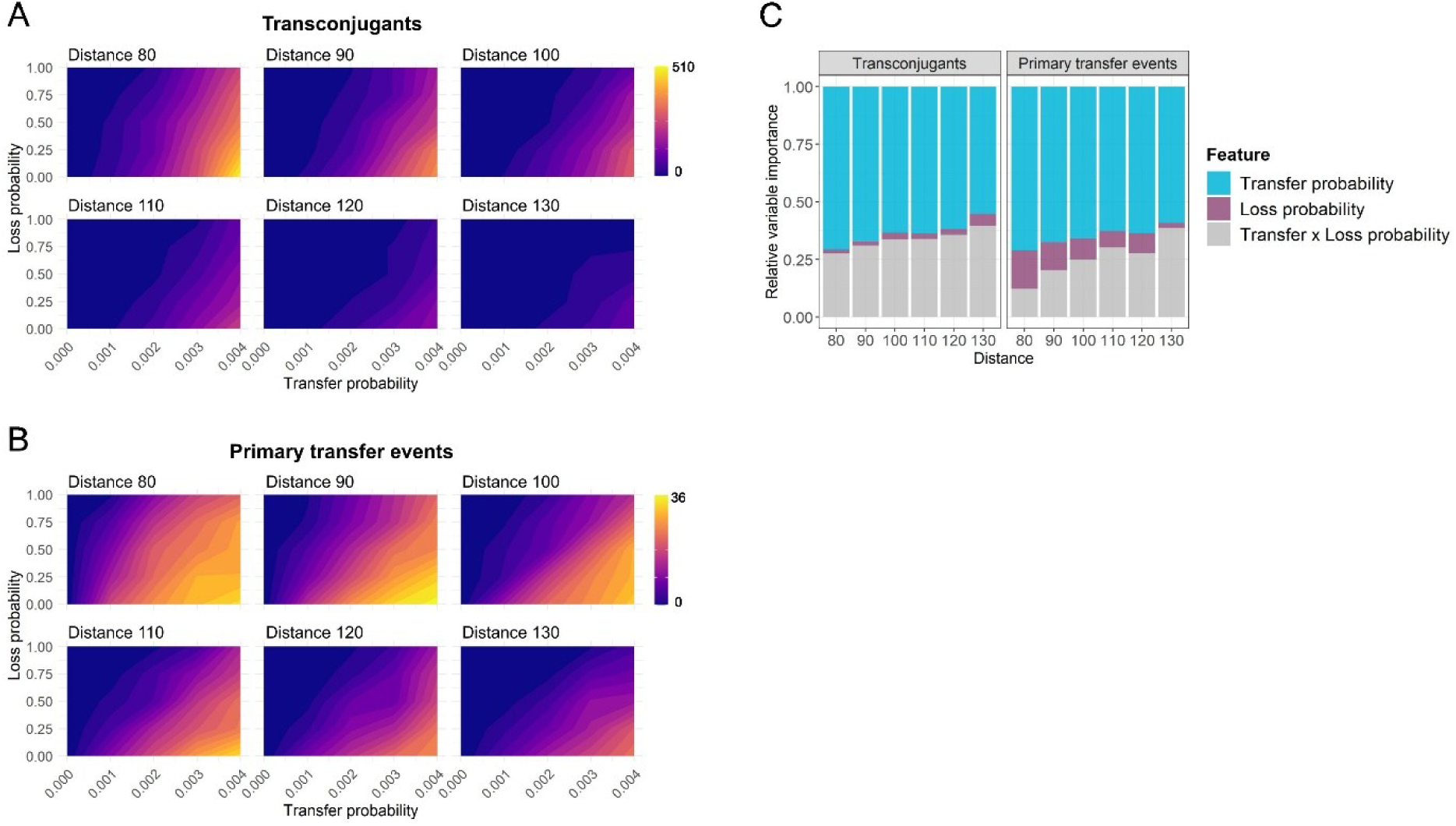
Interactive effects of plasmid transfer and loss probabilities with the initial distance between colonies on plasmid spread upon colony collision. Contour plots showing the interaction between the plasmid transfer probability, plasmid loss probability, and initial distance between inocula on (A) the total number of transconjugant cells, and (B) the total number of primary plasmid transfer events after collision of a plasmid donor and a potential recipient colony using an individual-based model. (C) Relative importance of the plasmid transfer and loss probabilities on plasmid spread as a function of the initial distance between inocula (n = 10). Individual-based simulations where the initial inoculum for the plasmid donor colony consisted of a 1:1 mixture of plasmid-carrying and -free cells, and the inoculum for the potential recipient colony consisted of a single population of plasmid-free cells (grey) which can receive the plasmid upon collision and become transconjugant cells (blue) (see Methods for full details).

For the total number of primary plasmid transfer events, the relative importance of the plasmid loss probability decreased from 16.7% at an initial distance of 80 (two-way ANOVA test; F*_1,250_* = 132.97, *P* < 2.2 x 10^-16^, n = 10) to 2.2% at an initial distance of 130 (two-way ANOVA test; F*_1,250_* = 227.35, *P* < 2.2 x 10^-16^, n = 10), while the relative importance of the plasmid transfer probability decreased from 71.0% at a distance of 80 (two-way ANOVA test; F*_1,250_* = 890.78, *P* < 2.2 x 10^-16^, n = 10) to 59.1% at an initial distance of 130 (two-way ANOVA test; F*_1,250_* = 385.62, *P* < 2.2 x 10^-16^, n = 10) (Fig. 3C). For the number of secondary plasmid transfer events within the potential recipient colony, we found that the relative importance of the plasmid loss probability increased and that of the plasmid transfer probability decreased as observed for the accumulation of transconjugant cells (Supplementary Fig. 8). This reveals that the initial spatial distribution of discrete colonies across a surface determines the effects of intrinsic drivers of plasmid dynamics.

### Local spatial intermixing determines plasmid spread upon colony collision

We hypothesized that the spatial organization of individual cells within the growing colonies would also be an important determinant of plasmid spread upon colony collision. As the initial distance between the colonies increases, we expect plasmid-carrying and -free cells within the plasmid donor colony to become increasingly spatially segregated due to longer growth times (as reported in [16, 17]). This, in turn, should reduce the extent of plasmid transfer both within the potential recipient colony during its growth (intraspecific secondary plasmid transfer) and between colonies after their collision (interspecific primary plasmid transfer). We used our individual-based modelling framework to simulate the collision boundary as two “cell walls”, where the plasmid donor cell wall consisted of plasmid-carrying (magenta) and -free (green) cells and the recipient cell wall consisted of a different population of plasmid-free (grey) cells (Fig. 4A). We varied the initial spatial intermixing of cells within the plasmid donor cell wall by either placing the plasmid-carrying and-free cells in two discrete patches (low spatial intermixing) or by sequentially alternating a cell of one type next to a cell of the other (high spatial intermixing) (Fig. 4A). After the cell walls collide, a single circular colony forms where the left and right hemispheres contain information regarding plasmid spread from the plasmid donor to the potential recipient cell wall. This simulation design allowed us to precisely control the spatial intermixing and organization of individual cells immediately before the point of collision and to investigate mechanisms occurring along the collision boundary.

**Figure 4.**
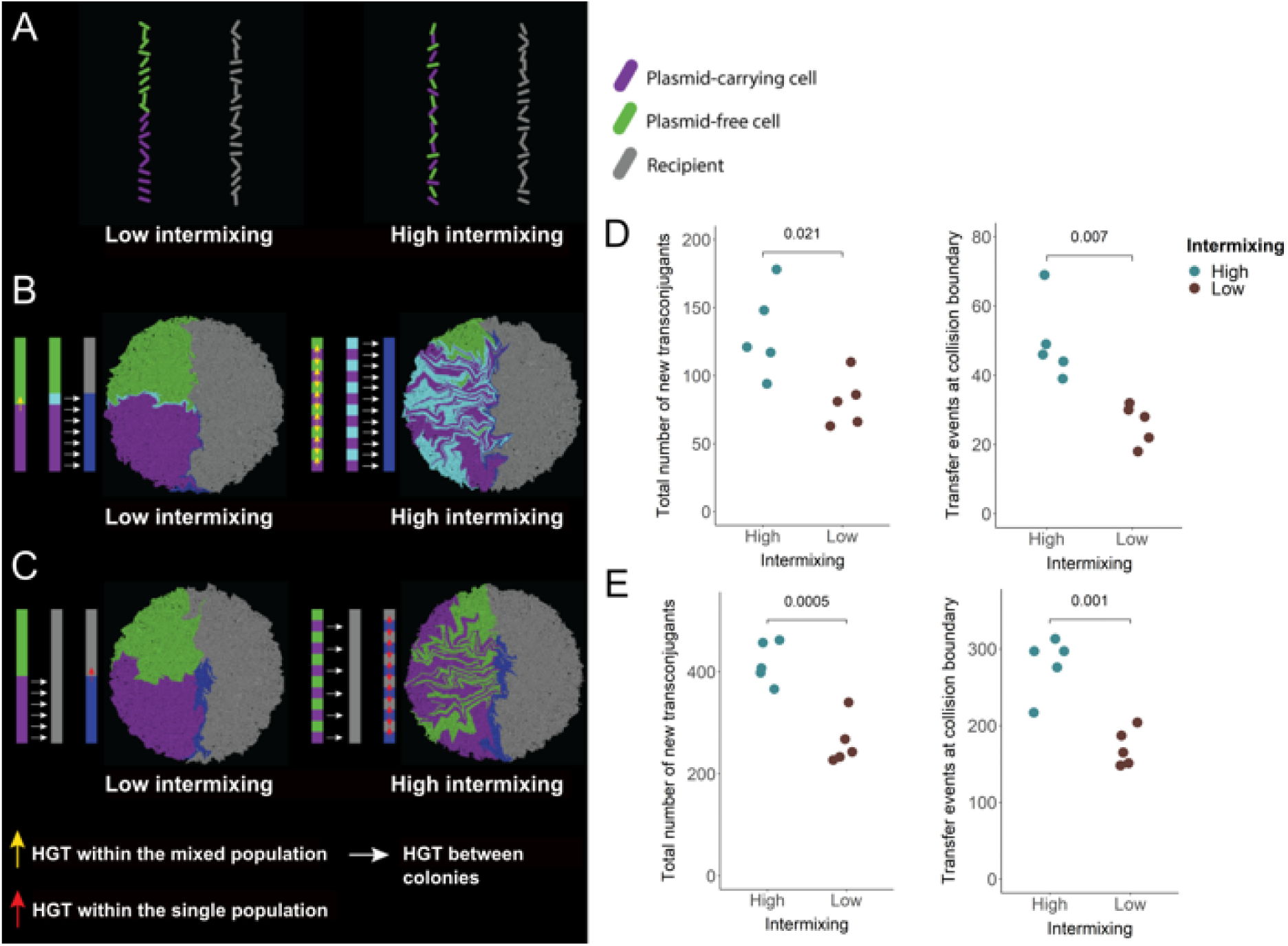
Effect of spatial intermixing of plasmid-carrying and -free cells along the collision boundary on plasmid spread. (A) Initial positions of cells along two cell walls that eventually collide. Plasmid-carrying (magenta) and -free (green) cells within the plasmid donor cell wall contain either two discrete patches (low intermixing) or alternate in cell-type (high intermixing). Potential recipient cells are grey. Cells are randomly rotationally oriented along the x-y plane. (B) Schematic figures of the expected effects are on the left and the simulation outputs are on the right. Transconjugant cells in the potential recipient cell wall are blue. (C) Simulations testing the effect of intermixing of plasmid -carrying and - free cells on plasmid spread. (D and E) Quantification of the (D) total number of transconjugant cells formed in the potential recipient hemisphere (blue) and (E) the total number of primary interspecific plasmid transfer events that occurred along the collision boundary for the simulations shown in (B) and (C), respectively. Simulation images correspond to the final time step. Each data point is for an independent simulation (n = 5). *P*-values on the top of (D) and (E) are for two-sample two-sided Welch t-tests.

We first tested the expectation that the extent of spatial intermixing will determine the total number of possible plasmid transfer events from the plasmid donor cell wall into the potential recipient cell wall by modulating the efficacy of interspecific plasmid transfer. To test this, we allowed plasmid transfer to occur within the plasmid donor cell wall and from the plasmid donor to the potential recipient cell wall but not within the potential recipient cell wall (i.e., we prevented intraspecific plasmid transfer within the potential recipient cell wall) (Fig. 4B). For collisions where plasmid-carrying cells (magenta) are highly intermixed with plasmid-free cells (green) in the plasmid donor cell wall, we found that most of the plasmid-free cells (green) become plasmid-carrying cells (cyan) due to intraspecific plasmid transfer, increasing the maximal plasmid load along the collision boundary (one-way ANOVA test; *F_1,2487_* = 1120, *P* = 2.0 x 10^-16^, n = 5) (Fig. 4B). Moreover, for high intermixing, we found that both the total number of new transconjugant cells (blue) and the number of interspecific plasmid transfer events along the collision boundary are higher than for low intermixing (two-sample two-sided Welch test; *P_Transconjugants_* = 0.021, *P_Transfer events_* = 0.007, n = 5) (Fig. 4D).

We next tested the expectation that the spatial intermixing of the plasmid-carrying and -free cells within the plasmid donor cell wall along the collision boundary will affect the spatial intermixing of transconjugant cells (blue) and plasmid-free (grey) cells in the potential recipient cell wall. Spatial features of the plasmid donor cell wall are acquired by the potential recipient cell wall upon collision and drive plasmid spread within the potential recipient cell wall by again modulating the efficiency of plasmid transfer (Fig. 4C). To test this, we allowed for plasmid transfer to occur between the plasmid donor and potential recipient cell walls and within the potential recipient cell wall, but not within the plasmid donor cell wall (i.e., we prevented intraspecific plasmid transfer within the plasmid donor cell wall). For collisions where plasmid-carrying cells (magenta) are highly intermixed with plasmid-free cells (green), we again found that both the total number of new transconjugant cells (blue) and the number of interspecific plasmid transfer events along the collision boundary are significantly higher (two-sample two-sided Welch test; *P* = 0.0005, *P* = 0.001, n = 5) (Fig. 4E). We then quantified the total number of plasmid transfer events that occurred within the potential recipient hemisphere and found that there were significantly more events for high intermixing (138 ± 21) compared to low intermixing (91 ± 28) (two-sample two-sided Welch test; *P* = 0.018, n = 5). Indeed, over one-third of plasmid spread within the potential recipient hemisphere was due to plasmid transfer between cells of the same cell-type (i.e. intraspecific transfer, 33.7 ± 5.1%).

### Experimental quantification of plasmid dynamics during bacterial colony growth and collision

To experimentally test the modelling results, we developed an experimental system that allows us grow bacterial colonies across nutrient rich surfaces, collide them into each other, and quantify plasmid dynamics during growth and collision. Briefly, we grew a plasmid donor colony containing two strains of *P. stutzeri*, where one carries pAR145 while the other does not, until it physically collided with a plasmid-free *E. coli* potential recipient colony. We first quantified pAR145 dynamics within the plasmid donor colony prior to its collision with the potential recipient colony. We expected the processes of pAR145 transfer and loss during the growth of the plasmid donor colony to affect the transfer of pAR145 into the potential recipient colony, as observed in the simulations (Fig. 1A,F and Fig. 2A). We found that pAR145 was nearly purged from the periphery of the plasmid donor colony after growing approximately 300 µm in the radial direction (corresponding to the radial interval between 1700-2000 µm) (Fig. 5A,B), which is the accumulated consequence of pAR145 loss and the faster growth of pAR145-free cells in the absence of chloramphenicol. Note that pAR145 was also readily transferred between plasmid-carrying and -free cells of *P. stutzeri* within the plasmid donor colony (see cyan areas in Fig. 5A). In our system, this radial interval between 1700-2000 µm determines the space/time where the interspecific transfer of pAR145 to *E. coli* cells within the potential recipient colony is likely to occur upon collision with the plasmid donor colony, as it determines the space/time when the plasmid is present along the periphery of the plasmid donor colony.

**Figure 5.**
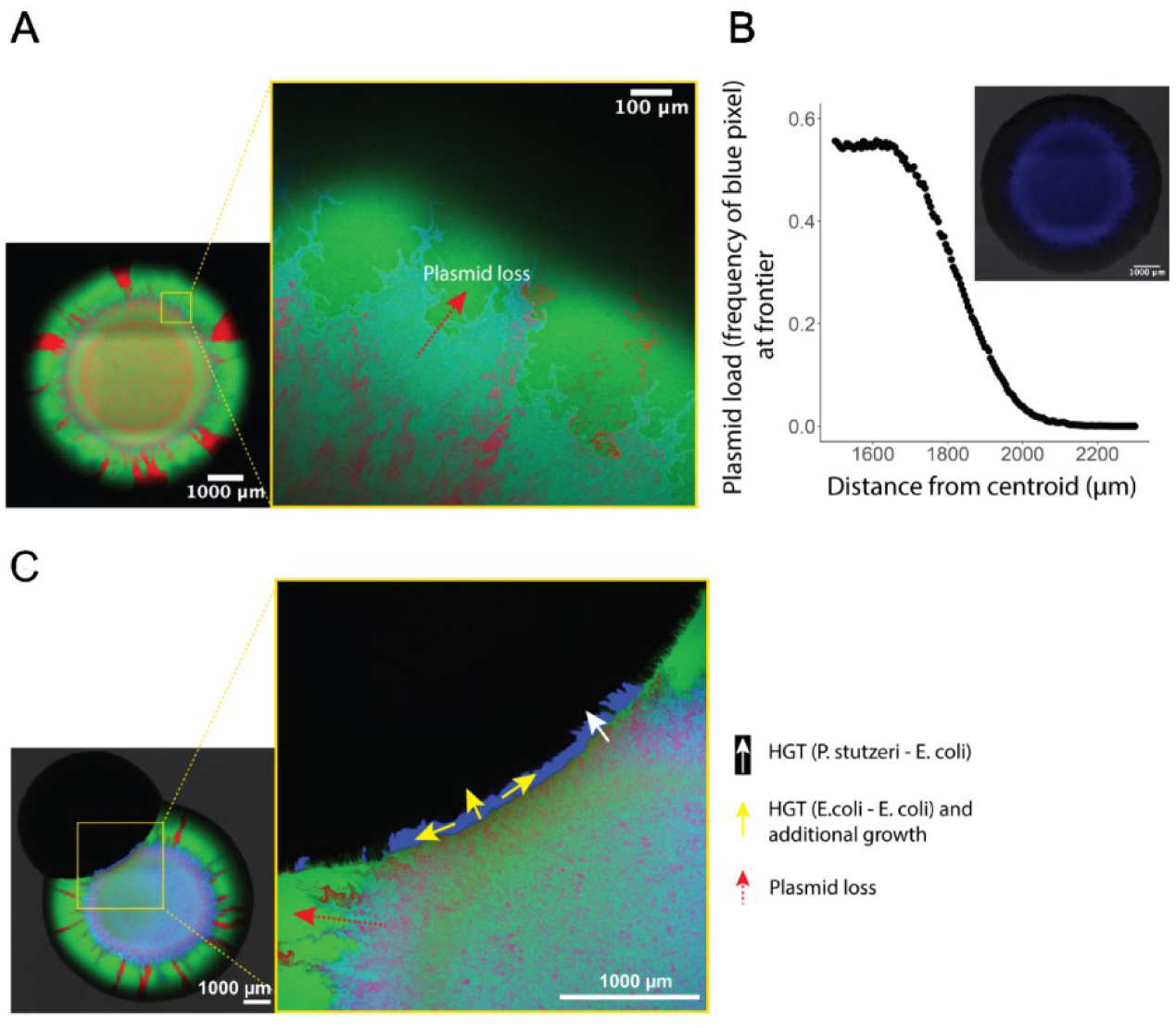
**Plasmid dynamics in plasmid donor colonies and interspecific transfer to potential recipient colonies upon collision. (**A) Representative microscopy image of the *P. stutzeri* plasmid donor colony consisting of two isogenic strains, where one strain contains pAR145 and expresses red fluorescent protein while the other is plasmid-free and expresses green fluorescent protein. If pAR145 transfers into the strain expressing green fluorescent protein, then it appears cyan. (B) Quantification of pAR145 abundance (blue pixels) during growth of the plasmid donor colony shown in (A). (C) Representative microscopy image of a collision between a *P. stutzeri* plasmid donor colony and a potential *E. coli* recipient colony. Note the formation of a blue patch located along the collision boundary, which corresponds to *E. coli* cells that acquired pAR145 via interspecific transfer from pAR145-carrying *P. stutzeri* cells.

### pAR145 transfer between colliding colonies depends on the initial distance between inocula

We next experimentally tested whether physical collisions between the *P. stutzeri* plasmid donor and the *E. coli* potential recipient colonies can promote interspecific pAR145 transfer into the potential recipient colony. To accomplish this, we placed a 1 µL droplet of the *P. stutzeri* inoculum at ca. 3.0 mm from the *E. coli* inoculum. After 96h of incubation, the plasmid donor and potential recipient colonies had grown and physically collided into each other. As predicted with our simulations, a new genotype formed along the collision boundary (Fig. 5C), which consisted of initially non-fluorescent *E. coli* potential recipient cells that obtained pAR145 from *P. stutzeri* plasmid donor cells upon colony collision. A closer evaluation of the collision boundary revealed that the newly formed *E. coli* transconjugant cells contiguously extended across an approximately 2 mm long boundary and uniformly protruded ca. 80 µm into the *E. coli* potential recipient colony (Fig. 5C).

After verifying that pAR145 can transfer between *P. stutzeri* plasmid donor and *E. coli* potential recipient colonies upon their physical collision, we next investigated the effect of the initial distance between inocula on this process (as done via simulation in Fig. 2). Following the simulation results, we hypothesized that the initial distance between the plasmid donor and potential recipient colonies would determine the extent of interspecific pAR145 transfer, where larger distances increase the time for pAR145 loss to occur from the plasmid donor colony prior to its collision with the potential recipient colony, and thus reduce pAR145 spread. To test this hypothesis, we used robotics to deposit the *P. stutzeri* plasmid donor and *E. coli* potential recipient inocula at precisely defined distances from each other and allowed them to develop into colonies that grew and eventually collided into each other. We found that collisions between colonies initially inoculated at closer distances led to larger amounts of new *E. coli* transconjugant cells upon collision than did colonies initially inoculated at longer distances (one-way ANOVA test; *F_1,28_* = 28.95, *P* = 9.8 x 10^-6^, n = 5) (Fig. 6A). The spatial range in which new *E. coli* transconjugant cells were formed were for initial distances between 2.8-3.3 mm, although the amount of new *E. coli* transconjugant cells decreased monotonically within this range (Spearman rank correlation test; rho = -0.793, *P* = 8.6 x 10^-5^, N = 30) (Fig. 6B).

**Figure 6.**
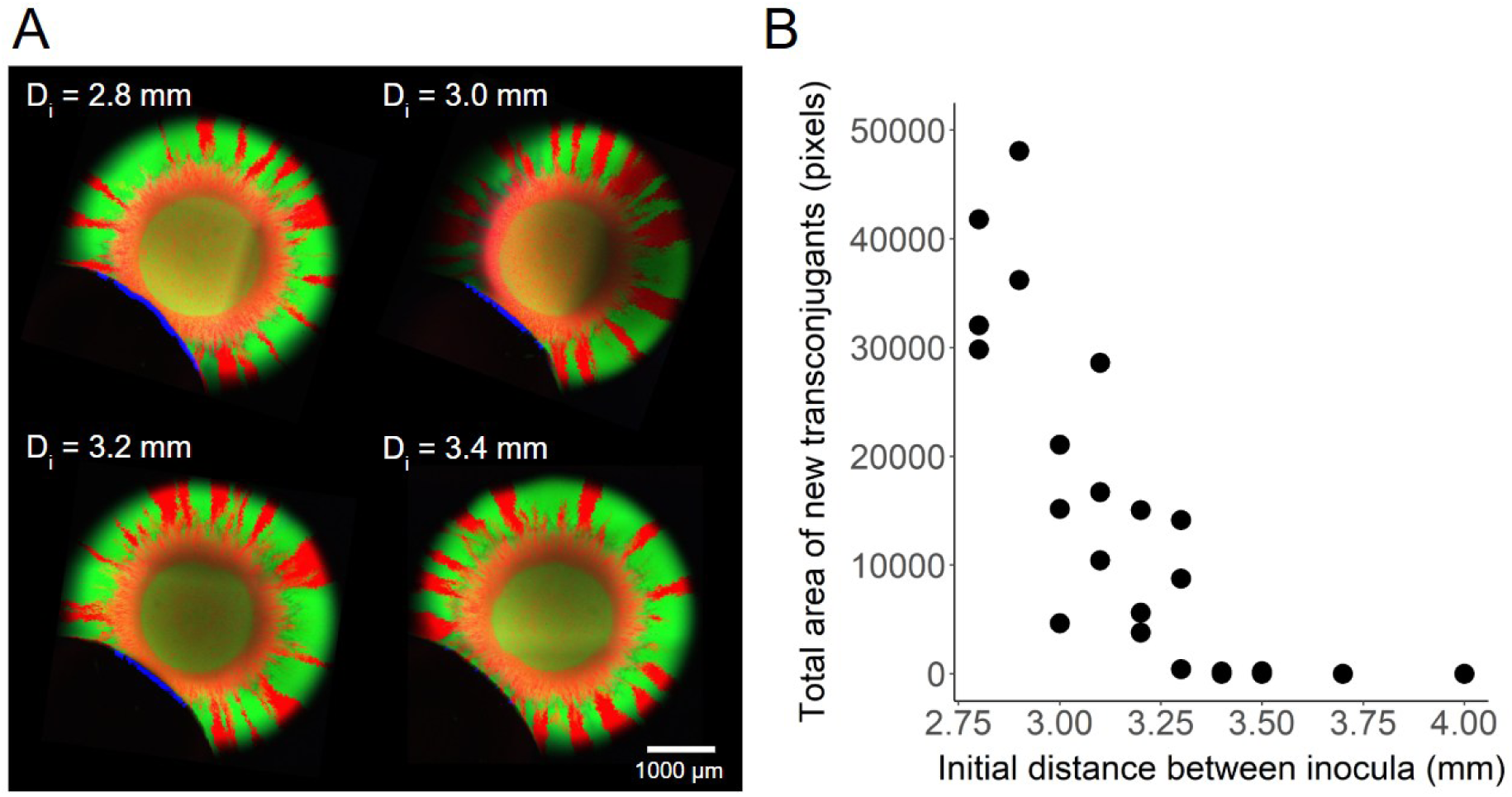
Initial distance between bacterial inocula determines the extent of interspecific pAR145 transfer upon colony collision. (A) Representative microscopy images of experimental collisions between *P. stutzeri* plasmid donor and *E. coli* potential recipient colonies. Numbers indicate the distances between the centroids of the inocula. Interspecific pAR145 transfer occurred along the collision boundaries and generated blue patches consisting of *E. coli* transconjugant cells carrying pAR145. Note that due to overexposure of the green and red channels, mixed populations of *P. stutzeri* do not display visible cyan signals even when carrying pAR145. *E. coli* does not express green or red fluorescent protein and thus appears blue when carrying pAR145. (B) Quantification of the absolute area of *E. coli* transconjugant cells as a function of the distance between the inocula. Each datapoint is for an independent biological replicate (n = 3) at the specified initial distance D_i_ (note that some datapoints are overlapping and thus appear as one).

## Discussion

Understanding the determinants of plasmid transfer both within and between surface-associated microbial biomass units is important for predicting the spread of antibiotic resistance and other plasmid-encoded traits, especially in environments where biomass is sparsely distributed. In this study, we showed that plasmid-encoded antibiotic resistance can readily transfer between spatially discrete microbial colonies during their growth and physical collision with each other (Fig. 5C). Our individual-based modelling simulations demonstrate that interactions between intrinsic properties of the plasmid-host system (plasmid transfer and loss probabilities, Fig. 1) and spatial factors (distances between biomass units, Fig. 2) are the primary drivers of plasmid spread. We combined controlled experiments to validate our simulations while also extending our simulations to test the generality of our findings across a broad range of plasmid transfer and loss probabilities and spatial conditions.

We further found that the importance of plasmid transfer and loss probabilities in determining plasmid spread was contingent on the initial distance between biomass units (Fig. 3). In the absence of antibiotic selection, there is a period during which plasmid loss upon cell division and selection for plasmid-free cells decreases the opportunities for interspecific plasmid transfer into adjacent colonies. Both in our experiments and simulations, this is evident as a decaying relationship between the extent of new transconjugant cells in the recipient colony and the initial distance between the colonies. Our analyses specifically determined that even if there are high plasmid transfer probabilities during this period, plasmid loss probabilities can partially offset these effects and become a sizeable determinant of plasmid spread into adjacent biomass units (Fig. 2G,H). The segregation control system of a particular plasmid will have a large impact on its temporal persistence [41, 42], and the cost of the plasmid will also determine how rapidly plasmid-free cells dominate the biomass periphery due to their higher intrinsic growth rate [43]. There are many cases in which plasmids can persist within the plasmid donor population even in the absence of positive selection due to compensatory mutations that reduce or eliminate the cost of plasmid carriage [44], high conjugation rates [20], or phage predation [45]. In such situations, the proliferation of plasmid free segregants would be substantially reduced, and we would therefore expect interspecific transfer to be independent of the initial distance between the colonies, as we would not expect a reduction in plasmid load to occur prior to collision. This means that the taxonomic or genotypic composition of a biomass unit and the biology of the antibiotic resistance-encoding plasmid will generally have large impacts on plasmid spread between adjacent biomass units [46].

We found that the spatial intermixing of plasmid-carrying and -free cells along the periphery of a plasmid donor colony affects both the total plasmid load and the degree of intermixing of the plasmid-free and -carrying cells in a potential recipient colony. More intermixed populations lead to a larger number of cell-cell contacts between distinct cell-types (in this case plasmid-free and -carrying types). For a contact-dependent process such as plasmid conjugation, this leads to a larger number of possible non-redundant plasmid transfer events (i.e., transfer events from plasmid donor to potential recipient cells as opposed to transfer events from plasmid donor to adjacent plasmid donor cells), which increases plasmid spread [17]. This explains why we observed a linear increase in the number of new transconjugant cells upon colony collision but a decelerating increase in the number of primary plasmid transfer events between colonies (Fig. 1E,J). After all the potential recipient cells in contact with a plasmid donor cell receive the plasmid along the collision boundary, there were no more plasmid-free cells for additional plasmid transfer, which explains the deceleration in the number of primary transfer events observed in our simulations.

In light of our findings, we expect that processes leading to higher spatial intermixing or spatial proximity between populations will increase plasmid spread. For example, in the mouse gut, the spread of AR-encoding plasmids is maximized with frequent contact between persisting plasmid donors and invading plasmid-free enteric pathogens [47, 48]. Frequent physical disturbances increase the number of new donor-recipient contacts by reshuffling the spatial positionings of cells and consequently promoting more extensive plasmid transfer [14]. Upon physical contact between biomass units, however, we confirmed previous experimental and theoretical work that transfer is more likely to occur at the biomass periphery [13], as shown by the narrow extent of the new *E. coli* transconjugant cells along the collision boundary. Seoane et al. [49] reported similar results, where there was interspecific plasmid transfer between small cell colonies, but plasmid invasion was limited by the inactivity of cells and the physical compression towards the colony center. This localized interspecific transfer can still be relevant for the maintenance of unstable plasmids in habitats where disturbances or environmental gradients promote the dispersal and regrowth of biomass associated, for example, with the mammalian gut, plant structures, or surfaces in aquatic systems (e.g. for gradients in oxygen concentrations [50]).

The link between plasmid dynamics within and between biomass units sets a basis for understanding antibiotic resistance spread at the metacommunity level, which better resembles processes occurring in natural systems such as in sparse biofilms or among collections of micro-colonies. The persistence of antibiotic resistance in microbial communities even in the absence of antibiotic pressure could be explained by spatial factors such as the spatial intermixing between cells that operate both within and between biomass units. In spatially structured systems such as on medical devices or dental plaque, this conceptualization might be of interest because the spread of plasmid-encoded antibiotic resistance could be simulated based on pre-existing models from metacommunity theory [51, 52]. Source-sink dynamics are a way by which a stable plasmid donor ensures the plasmid is maintained in unstable plasmid recipients by frequent transfer [19]. Our study suggests that depending on the physical proximity between sources and sinks of plasmid transfer, frequent collisions between biomass units can maintain plasmids via transfer even when the plasmid is unstable in all community members. This implies that in a metacommunity context, the entire metacommunity will have access to the plasmid under source-sink dynamics, provided there is frequent physical contact between biomass units.

The relative simplicity of our experimental system and modelling approach also has several limitations. Our simplified consortia can effectively minimize confounding factors and enable us to draw strong conclusions, but this comes with the cost of not being able to capture all the complex interactions and processes within collections of biomass units such as in sparse multi-species biofilms. However, the processes we describe here should nevertheless be important, as different cell-types will persist at the biomass periphery and be susceptible to plasmid transfer. Second, we have not tested the dependency of plasmid transfer on the cell-type composition of the adjacent colliding biomass units. Some cell-types can produce a thick layer of exopolysaccharides that can prevent cell-cell contacts even when there are compressive physical forces at play [53]. Also, some cell-types might not have compatible conjugation machineries to establish pili junctions required for effective transfer [54], while some biofilms will prevent collisions in the first place via chemical signaling [55]. The generalizability of our findings could be confirmed with studies that implement our simplified experimental system using diverse sets of strains from multiple taxonomic groups with varying traits. There are also multiple factors we could not address that will determine the extent of this spread into recipient biomass. First, physical forces might push new transconjugants towards the collision boundary, uplifting them and creating a vertical rather than a horizontal expansion [51]. Second, the metabolic state of cells closer to the center of the biomass will also determine their ability to capture and transfer the plasmid. Here nutrient availability, interspecies interactions, and abiotic stressors might play an important role in determining such cellular activity. Finally, the presence of even subinhibitory concentrations of the antibiotic, to which a plasmid confers resistance, can create a positive selective pressure that promotes further transfer.

### Conclusions

Our study reveals how the interplay between plasmid dynamics and microbial spatial organization determine the spread of plasmid-encoded antibiotic resistance, offering opportunities to anticipate and manage this global challenge. By linking plasmid transfer and loss probabilities to spatial factors such as physical distances between microbial biomass units and intermixing between distinct populations, we establish quantitative relationships that can be integrated into predictive models of resistance dissemination across natural and clinical environments. These insights can facilitate the design of strategies that limit resistance spread on surfaces [56], for example via surface engineering [57], targeted treatment of discrete plasmid-carrying populations [58], or accounting for the 3D structure of biofilms [59]. Ultimately, our framework moves toward a more mechanistic and predictive understanding of when and where antibiotic resistance will persist within spatially structured microbial ecosystems.

## Methods

### Bacterial strains and plasmid

The plasmid donor colonies consisted of two genetically engineered isogenic mutants of the bacterium *Pseudomonas stutzeri* A1601, whose genetic modifications and growth traits were established previously [60, 61]. Each strain is genetically identical to the other except for carrying a different isopropyl β-D-1-thiogalactopyranoside (IPGT)-inducible fluorescent protein-encoding gene located on the chromosome (*egfp* encoding for green fluorescent protein or *echerry* encoding for red fluorescent protein) (Supplementary Table 5). This enables us to distinguish and quantify the different strains when grown together in a spatially explicit manner. We transformed the strain carrying the *echerry* gene with the low copy number plasmid pAR145 (pSU2007 *aph*::*cat*-P_A1/04/03_-*cfp*^∗^-T_0_, described in [62]). pAR145 is a derivative of plasmid R388 and encodes for chloramphenicol resistance and an IPTG-inducible *ecfp* gene (encoding for cyan fluorescent protein). The potential recipient strain was *E. coli* DH5α [F2 *supE44 lacU169 (w80lacZDM15) hsdR17 recA1 endA1 gyrA96thi-1 r elA1*] [63].

### Quantification of plasmid cost

We used a previously described colony collision assay [64] to measure the cost of carrying pAR145 in terms of its effect on growth rate in the absence of positive selection. We grew the pAR145-carrying and -free *P. stutzeri* strains in isolation and adjusted the OD_600_ of each overnight culture to 2.0 with 0.89% (w/v) sodium chloride solution. We then placed a 1 µL drop of the pAR145 donor mixture and another 1 µL drop of the potential recipient 3 mm apart from each other on replicated lysogeny broth (LB) agar plates (1.5% agar). We incubated the plates at room temperature for 96h, which allowed the colonies to grow and eventually collide into each other. We estimated the relative growth rates of the pAR145 donor and potential recipient strains using the curvature of the arc of the collision boundary between the colonies and the radii of the colonies using the following equation,

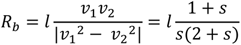

where *l* is the distance between two colonies, *s* is the cost of carrying pAR145, *v*_1_ is the expansion velocity of the pAR145-free colony, and *v*_2_ is the expansion velocity of the pAR145-carrying colony. *R*_*b*_ is the radius of the circle generated by the arc at the collision boundary, which together with the mentioned parameters allows the derivation of *s*. We quantified *R*_*b*_ and *l* for four biological replicates using Adobe Illustrator (v27.0.1). See Supplementary Table 1 for detailed data. These calculations allowed us to set realistic parameters for our individual-based modelling (see below).

### Individual-based modelling of colony collisions

We performed individual-based modelling of colony growth and collision using CellModeller 4.3, a Python-based computational framework designed to model the growth and division of multicellular systems [40]. We ran all simulations on the ETH Euler high-performance computing cluster using the Slurm workload manager for batch job submissions. We modelled individual bacteria as three-dimensional rod-shaped capsules using the CLBacterium biophysical module with growth restricted to a monolayer. We initialised cells to have a length of 3.5 units and allowed them to elongate along their long axis at a rate set by their growth rate parameter with growth drag coefficient gamma = 20. For each newly born cell, we defined an individual division threshold by adding a Gaussian-distributed increment (mean 2.0 and standard deviation 0.45) to its length at birth. Cells divided when their size exceeded this threshold, producing two daughter cells of approximately half the parent cell’s division length and each assigned a new division threshold. This constant added-size (length) can accurately model bacterial division and cell size homeostasis [65]. We ran each simulation until reaching a total population size of 20,000 cells.

To study plasmid dynamics during colony expansion and collision, we simulated two spatially separated biomass units composed of strains with distinct backgrounds, which we refer to as the plasmid donor and potential recipient colonies. We then allowed them to grow and physically collide into each other. We encoded each combination of colony background (plasmid donor or potential recipient) and plasmid state (plasmid-free or -carrying) as a discrete cell-type and updated these labels whenever plasmid transfer or loss occurred. For the simulations reported in Figs. 1-3, we initialized both colonies as circles (radius 40 units) with their centers d units apart (d = 80, 90, 100, 110, 120, 130). The donor colony contained 200 cells placed at random positions within the circle in a 1:1 mixture of plasmid-free (green) and -carrying (magenta) cells. The potential recipient colony contained 200 plasmid-free cells (grey) placed at random positions within the second circle on the same xy-plane. We set the growth rate = 1.0 for plasmid-free in the plasmid donor-colony, 0.83 for plasmid-carrying cells in the plasmid donor colony, and 0.8 for both the plasmid-free and - carrying cells in the potential recipient colony, matching our experimentally measured plasmid costs (reported in [17]). Thus, plasmid carriage imposed a 17% growth-rate cost in the plasmid donor cells but no additional cost in the potential recipient cells.

We extended the model to incorporate plasmid transfer and loss. We simulated contact-dependent plasmid transfer using the neighbor lists returned from the CLBacterium (set by parameter ‘compNeighbours=True’) [17]. At each time step, a plasmid-free cell in contact with a plasmid-carrying cell switches to the corresponding plasmid-carrying cell-type with a defined probability (referred to as the plasmid transfer probability). We simulated plasmid loss during cell division. At each cell division, one of the two daughter cells is chosen at random and switches to a plasmid-free cell of the same cell-type with a defined probability (referred to as the plasmid loss probability). We assumed that plasmid-carrying cells in the potential recipient colony were stable (plasmid loss probability = 0) as we observed experimentally. We tracked all plasmid transfer and loss events using dedicated counters throughout each simulation. We performed 1500 unique simulations using the following full factorial design: Plasmid transfer probability (5 levels; 0-0.001-0.002-0.003-0.004) x plasmid loss probability (5 levels; 0-0.25-0.5-0.75-1.0) x distance between inocula (6 levels; 80-90-100-110-120-130) x 10 replicates.

To test the effect of spatial intermixing of plasmid-donor and -free cells on plasmid spread, we placed cells from the plasmid donor and potential recipient cell-types along two parallel cell walls. The plasmid donor wall contained a 1:1 mixture of plasmid-carrying and -free cell while the potential recipient wall contained only plasmid-free cells. We assigned the initial rotational orientation of each cell at random. In this setup, we considered two initial configurations for the donor boundary: a pattern with high intermixing (alternation of plasmid-carrying and plasmid-free cells) and a pattern with low intermixing (spatial segregation by type). We set the growth rate of all cell-types to be the same because the length and time scales are small when simulating collision boundaries and to isolate the effects of spatial positioning from differences in growth rates. We used the same values for all other settings used for the simulations, except that we removed the plasmid loss process for this short time scale.

### Colony collision experiments

We performed colony collision experiments where we inoculated a mixture of the *P. stutzeri* strains (the plasmid donor colony) and *E. coli* DH5α (the potential recipient colony) as single droplets onto replicated LB agar plates and allowed them to grow until they physically collide with each other. We initially grew all the strains separately in oxic liquid LB medium overnight at 37°C and set their optical densities at 600 nm (OD_600_) to 2. We next mixed the two *P. stutzeri* strains (plasmid-carrying and -free) at a 1:1 ratio (vol:vol). The *echerry*-expressing plasmid donor strain contained pAR145, which encodes for *ecfp*, and displayed the composite color magenta (cyan and blue). The *egfp*-expressing potential recipient strain only displays the color green. If the potential recipient strain receives pAR145, then it displays the color cyan. Using an Evo 200 robotic liquid handling system (Tecan, Männedorf, Switzerland), we deposited pairs of droplets of the *P. stutzeri* mixture and the *E. coli* strain (1 µl of each culture) at four discrete spatial positions on each LB agar plate adjusted to a pH of 7.5 and amended with 1 mM IPTG. We programmed the liquid handling system to deposit the *P. stutzeri* mixture and *E. coli* droplets at distances between droplet centroids of 2.80, 2.90, 3.00, 3.10, 3.20, 3.30, 3.40, 3.50, 3.70, and 4.00 mm with 3 replicates per distance. We incubated the plates for 96h at room temperature in oxic conditions.

### Bacterial colony imaging and quantitative image analysis

We imaged the colonies immediately after completion of the incubation period using a Leica TC5 SP5 II confocal microscope (Leica Microsystems, Wetzlar, Germany) with objectives 10x/0.3na (dry) and 63 x/1.4na (Wetzlar, Germany). We scanned the entire colony areas (*P. stutzeri* mixture and *E. coli*) by stitching together multiple frames of 1024 x 1024 pixels. We set the laser emission to 514 nm for the excitation of red fluorescent protein, 488 nm for the excitation of green fluorescent protein, and 458 nm for the excitation of cyan fluorescent protein. We analyzed the images in ImageJ; https://imagej.nih.gov/ij/) using FIJI plugins (v.2.1.0/1.53c; https://fiji.sc).

We quantified plasmid load (i.e., the total area of cyan fluorescent protein-expressing cells) using the “Sholl analysis” plugin [63] on the binarized image of the cyan channel after application of a noise-reduction threshold of 20 pixels. The “Sholl analysis” calculated the number of cyan pixels at 10 µm radial increments from the centroid to the colony edge. We initiated the analysis at 1500 µm from the colony centroid because this distance represents the edge of the inoculation droplet. We then applied the “Area to line” function of the “Overlay” plugin to register the number of cyan pixels for the areas corresponding to each 10 µm radial increment. We defined plasmid load as the total number of cyan pixels divided by the total number of pixels along each radius.

### Statistical analysis

We performed treatment comparisons using one-way ANOVA when we had sufficient data to test whether our datasets deviate from normality using the Shapiro-Wilk test. We performed treatment comparisons using two-sample two-sided Welch tests when we lacked sufficient data to test whether our datasets deviate from normality. We estimated the relative factor importance of transfer and loss probabilities on plasmid spread using two-way ANOVA with the interaction between these factors using the base R package *relaimpo* (v2.2-7). We tested for correlations between the initial distance between colliding colonies and the extent of plasmid transfer between colonies using Spearman rank correlation tests. We tested for linear relationships between simulated colony growth and plasmid spread using linear regression. We performed all statistics in R (v4.4.1, https://www.r-project.org/).

## Acknowledgments

We thank Dr. María Pilar Garcillán-Barcia (University of Cantabria) for providing plasmid pAR145 and *E. coli* DH5α. J.R. acknowledges funding from the Spanish Agency of Research (MicroDroughtPredict project: PID2023-151209NA-I00), the Swiss National Science Foundation (Early Postdoc Mobility grant: P2EZP3_199849) and the Generalitat de Catalunya (Beatriu de Pinós grant: 2022-BP-00050). M.M. acknowledges funding from the Swiss National Science Foundation awarded to M.M. (Ambizione grant: PZ00P3_180147). Y.M. acknowledges funding from the Swiss National Science Foundation awarded to D.R.J. (Individual project grants: 31003A_176101 and 310030_207471). D.R.J. acknowledges funding from an Eawag Discretionary Funds grant (category SEED).

## Author contributions

J.R., Y.M. and D.R.J. conceptualized the study and designed the experiments. J.R., Y.M., and J.S. performed the experiments with assistance from M.M. and D.A. Y.M., D.V. and C.R. performed the model simulations. J.R. and Y.M. analyzed the data. J.R. wrote the manuscript with input from all co-authors. J.R. and D.R.J. coordinated the study.

## Data availability

All data and code generated in this study will be deposited in the Eawag Research Data Institutional Repository (https://opendata.eawag.ch/) at the point of revision and will be made freely available to the public. All data and code are currently available at https://drive.google.com/drive/folders/1j7gIvDwmMp88zCvzqnTi-87d_PETSQww?usp=sharing.

