## Supplementary for "Plasmid transfer and loss probabilities interactively determine antibiotic resistance spread in surface-associated bacterial biomass"

### Supplementary materials

**Supplementary Table 1.** Quantification of relative growth rates of *P. stutzeri* A1601-*ech* with pAR145 and *P. stutzeri* A1601-*gfp*. $s$ is the selective advantage, or in our case the cost, that the plasmid bestows in the absence of chloramphenicol; $R_b$ is the radius of the circle generated by the arc at the boundary; $l$ is the distance between centroids of two colonies. $R_b$ and $l$ are expressed in μm.

| Replicate | $s$ | $R_b$ | $l$ |
|---|---|---|---|
| 1 | 0.834 | 207.552 | 268.231 |
| 2 | 0.84 | 230.751 | 298.679 |
| 3 | 0.827 | 230.756 | 295.293 |
| 4 | 0.83 | 230.742 | 295.799 |

**Supplementary Table 2.** Summary of the generalized linear model for testing the full effects of the plasmid transfer probability, plasmid loss probability, and initial distance between colonies on the total number of transconjugants and the number of primary plasmid transfer events after collision of a plasmid donor and a potential recipient colony. We performed a total of 1500 simulations. Full model formula: *Y ~ Transfer probability* + *Loss probability* + *Distance* + *Transfer:Loss* + *Transfer:Distance* + *Loss:Distance* + *Transfer:Loss:Distance* (n = 10).

| | Total number of transconjugant cells | | | Total number of primary transfer events | | |
|---|---|---|---|---|---|---|
| **Factor** | **Estimate** | ***t*** | ***P*** | **Estimate** | ***t*** | ***P*** |
| Transfer probability | 326300 | 34.29 | $< 2.2 \times 10^{-16}$ | 14340 | 13.83 | $< 2.2 \times 10^{-16}$ |
| Loss probability | 22 | 0.58 | 0.563 | -19.04 | -4.59 | $4.77 \times 10^{-6}$ |
| Distance | 0.93 | 4.24 | $2.34 \times 10^{-5}$ | -0.12 | -5.17 | $2.67 \times 10^{-7}$ |
| Transfer x Loss probability | -152300 | -9.80 | $< 2.2 \times 10^{-16}$ | 1559 | 0.92 | 0.357 |
| Distance x Transfer probability | -2294 | -25.64 | $< 2.2 \times 10^{-16}$ | -56.21 | -5.77 | $9.77 \times 10^{-9}$ |
| Distance x Loss probability | -0.09 | -0.25 | 0.800 | 0.13 | 3.46 | $5.59 \times 10^{-4}$ |
| Distance x Transfer x Loss probability | 945 | 6.47 | $1.34 \times 10^{-10}$ | -54.51 | -3.43 | $6.3 \times 10^{-4}$ |

**Supplementary Table 3.** Summary of the generalized linear model for testing the full effects of the plasmid transfer probability, plasmid loss probability, and initial distance between inocula on the total number of transconjugants and of primary plasmid transfer events after collision of a plasmid donor and a potential recipient colony. We performed a total of 1500 simulations and quantified the total number of transconjugant cells and the number of accumulated plasmid transfer events at a fixed simulation time point of 640. Full model formula: *Y ~ Transfer probability* + *Loss probability* + *Distance* + *Transfer:Loss* + *Transfer:Distance* + *Loss:Distance* + *Transfer:Loss:Distance* (n = 10).

| | **Total number of transconjugants** | | | **Total number of primary transfer events** | | |
|---|---|---|---|---|---|---|
| **Factor** | **Estimate** | ***t*** | ***P*** | **Estimate** | ***t*** | ***P*** |
| Transfer probability | 205100 | 30.30 | $< 2.2 \times 10^{-16}$ | 12610 | 13.55 | $< 2.2 \times 10^{-16}$ |
| Loss probability | 1.35 | 0.50 | 0.617 | -15.15 | -4.07 | $4.89 \times 10^{-5}$ |
| Distance | 0.58 | 3.73 | $1.2 \times 10^{-4}$ | -0.094 | -4.38 | $1.26 \times 10^{-5}$ |
| Transfer x Loss probability | -92670 | -8.41 | $< 2.2 \times 10^{-16}$ | 597.60 | 0.39 | 0.694 |
| Distance x Transfer probability | -1424 | -22.45 | $< 2.2 \times 10^{-16}$ | -50.43 | -5.77 | $9.80 \times 10^{-9}$ |
| Distance x Loss probability | -0.07 | -0.28 | 0.777 | 0.11 | 3.07 | 0.002 |
| Distance x Transfer x Loss probability | 566.70 | 5.47 | $5.21 \times 10^{-8}$ | -41.35 | -2.90 | 0.004 |

**Supplementary Table 4.** Summary of the generalized linear model for testing the full effects of the plasmid transfer probability, plasmid loss probability, and initial distance between inocula on the total number of secondary plasmid transfer events after collision of a plasmid donor and a potential recipient colony. We performed a total of 1500 simulations. Full model formula: *Y ~ Transfer probability* + *Loss probability* + *Distance* + *Transfer:Loss* + *Transfer:Distance* + *Loss:Distance* + *Transfer:Loss:Distance* (n = 10).

| | **Total number of secondary transfer events** | | |
|---|---|---|---|
| **Factor** | **Estimate** | ***t*** | ***P*** |
| Transfer probability | 228500 | 34.52 | $< 2.2 \times 10^{-16}$ |
| Loss probability | 37.55 | 1.42 | 0.156 |
| Distance | 0.86 | 5.61 | $2.40 \times 10^{-8}$ |
| Transfer x Loss probability | -115100 | -10.65 | $< 2.2 \times 10^{-16}$ |
| Distance x Transfer probability | -1694 | -27.23 | $< 2.2 \times 10^{-16}$ |
| Distance x Loss probability | -0.24 | -0.97 | 0.334 |
| Distance x Transfer x Loss probability | 791.10 | 7.79 | $1.29 \times 10^{-14}$ |

**Supplementary Table 5.** Specifications of the strains and plasmid used in this study.

| Strain | Relevant characteristics | Reference |
|---|---|---|
| *P. stutzeri* A1601-*egfp* | *P. stutzeri* A1501 (wild type) with Δ*comA* and mini-Tn7T-LAC-Gm-*egfp*; $Gm^R$, *egfp*$^+$ | 60,61 |
| *P. stutzeri* A1601-*ech* | *P. stutzeri* A1501 (wild type) with Δ*comA* and mini-Tn7T-LAC-Gm-*ech*; $Gm^R$, *ech*$^+$ | 60,61 |
| *E. coli* DH5α | F2 *supE44 lacU169 (w80lacZDM15) hsdR17 recA1 endA1 gyrA96thi-1 r elA1* | 62 |
| pAR145ecfp | pSU2007 *aph::cat*-PA1/04/03-*cfp*⁄-T0 | 62 |

A B

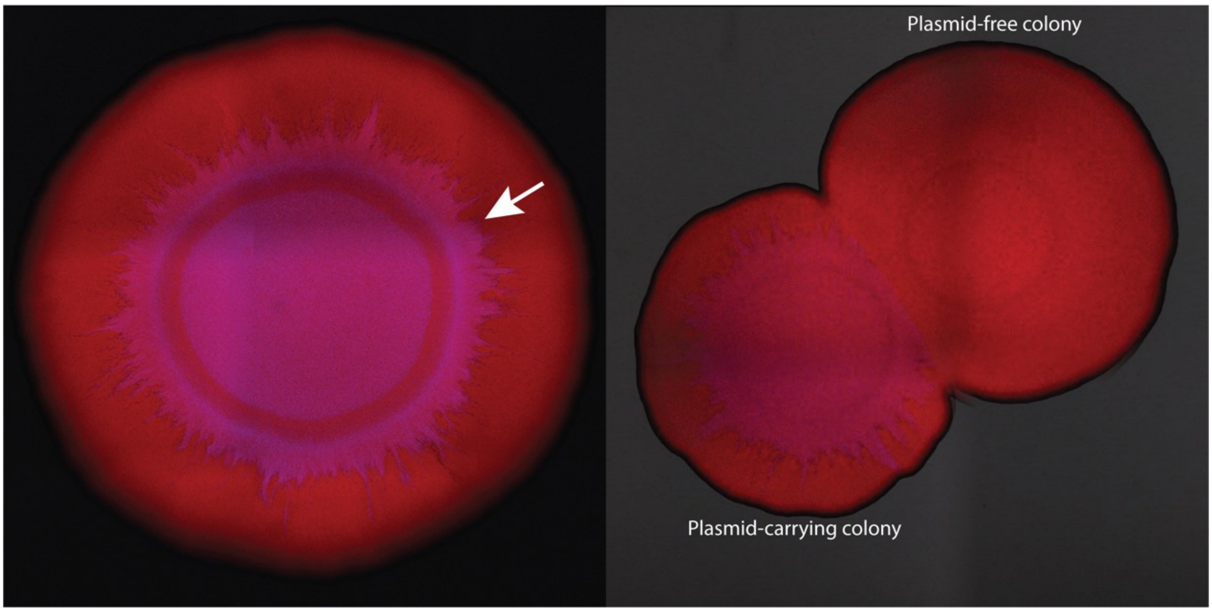


**Supplementary Figure 1.** Loss of pAR145 in the absence of chloramphenicol during colony growth. (A) We started the colony experiment with a single pAR145-carrying strain of *P. stutzeri* A1601 that encodes for *echerry* on its chromosome. It therefore initially displays both red fluorescence from the chromosome and blue fluorescence from pAR145 and appears the composite color magenta. The white arrow identifies a pAR145 loss event indicated by a change in color from magenta to red. (B) We performed the collision experiment between *P. stutzeri* A1601-*ech* carrying pAR145 (magenta) and *P. stutzeri* A1601-*ech* (red). We inoculated the two strains at the same time at a distance of approximately 3 mm away from each other. We took images after 96 hours of incubation at room temperature. The smaller size of the pAR145-carrying colony indicates a slower growth rate, demonstrating that carrying pAR145 is costly in the absence of chloramphenicol.

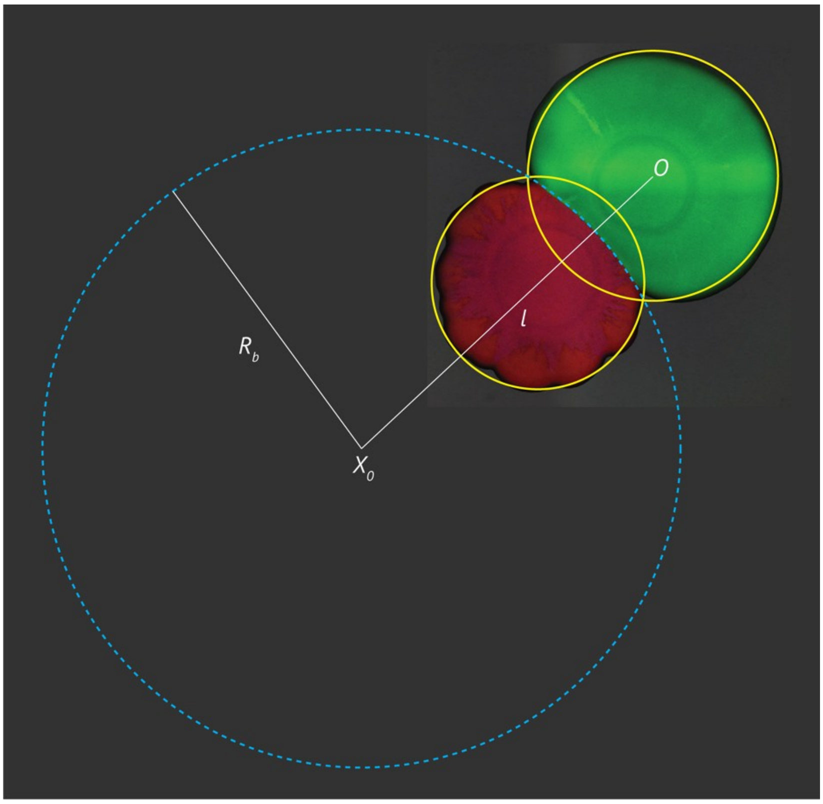


**Supplementary Figure 2**. Collision assay of *P. stutzeri* A1601-*ech* carrying pAR145 and *P. stutzeri* A1601-*gfp*. We drew the larger circle based on the arc generated between the two colonies. $X_0$ and *O* are the centroids of the derived circle and the green colony, respectively. $l$ is the distance between $X_0$ and *O*. $R_b$ is the radius of the derived circle. We use following equation to calculate the relative growth rate $s$.

$$R_b = l\frac{1+s}{s(2+s)}$$

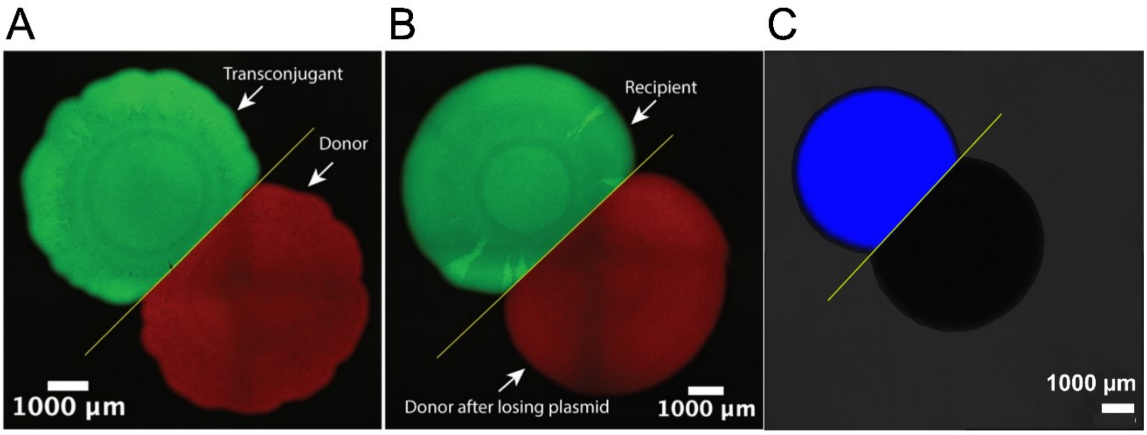


**Supplementary Figure 3.** Collision assays of *P. stutzeri* A1601-*gfp*, *P. stutzeri* A1601-*ech*, and *E. coli* DH5α either carrying or not carrying pAR145. The straight yellow lines at the collision interfaces indicate equal growth rates of the two strains. (A) Collision assay of *P.*

*stutzeri* A1601-*ech* carrying pAR145 (plasmid donor) and *P. stutzeri* A1601-*gfp* carrying pAR145. (B) Collision assay of *P. stutzeri* A1601-*ech* (plasmid donor after losing pAR145) and *P. stutzeri* A1601-*gfp*. (C) Collision assay of *E. coli* DH5α (black) and *E. coli* DH5α carrying pAR145 (blue).

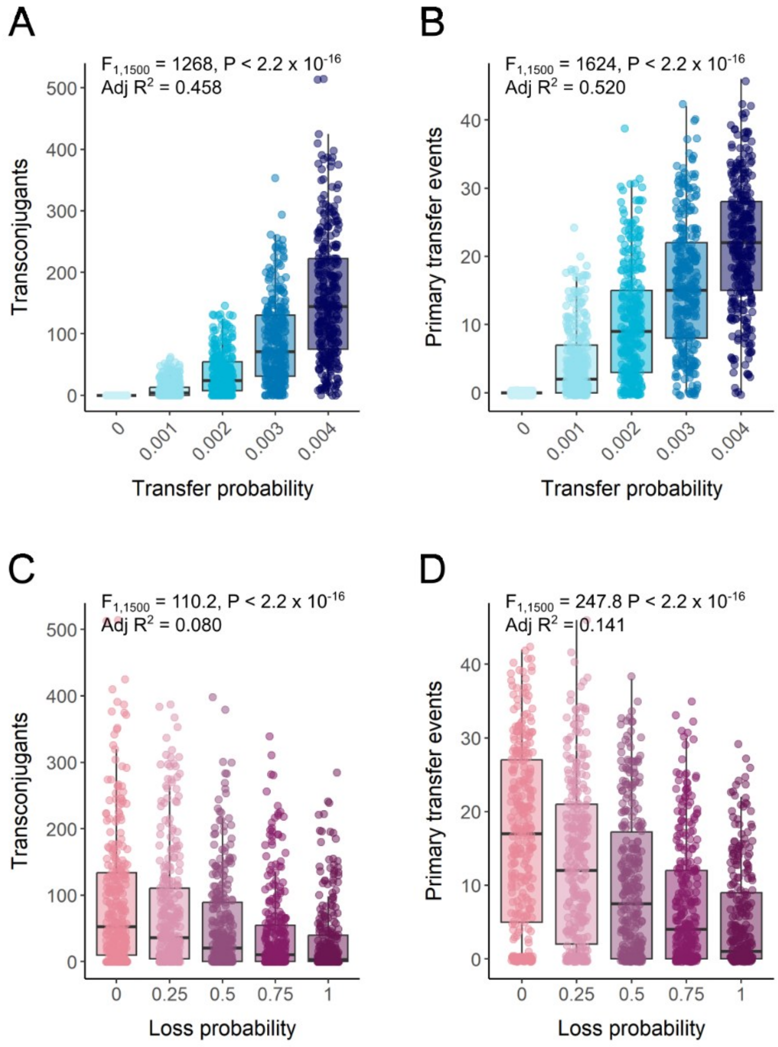


**Supplementary Figure 4.** Effect of plasmid transfer and loss probabilities on plasmid spread between colliding colonies at a fixed simulation time step of 640. (A) Total number of transconjugant cells (blues) as a function of the plasmid transfer probability. (B) Total number of primary plasmid transfer events between the plasmid donor and potential recipient colonies as a function of the plasmid transfer probability. (C) Total number of transconjugant cells (blue) as a function of the plasmid loss probability. (D) Total number of primary plasmid transfer events between the plasmid donor and potential recipient colonies as a function of the plasmid loss probability. Statistics are for one-way ANOVA tests with the (A, B) plasmid transfer probability or (C, D) plasmid loss probability as the sole explanatory variable (n = 10).

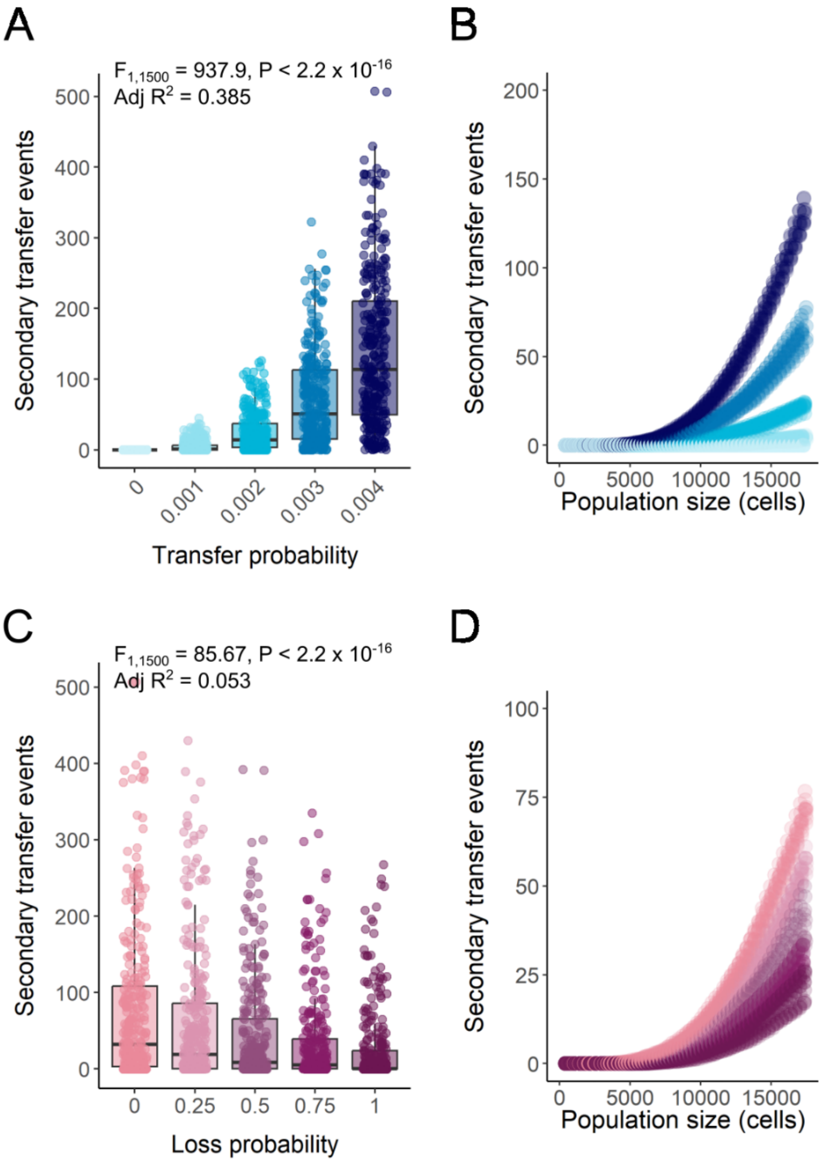


**Supplementary Figure 5.** Effect of plasmid transfer and loss probabilities on the total number of secondary plasmid transfer events within the potential recipient colony. (A) Total number of secondary plasmid transfer events at the end of the simulation as a function of the plasmid transfer probability. (B) Total number of transconjugant cells (blue) as a function of the total population size, colored by plasmid transfer probability as defined in (A). (C) Total number of secondary plasmid transfer events at the end of the simulation as a function of the plasmid loss probability. (D) Total number of secondary plasmid transfer events as a function of the total population size, colored by the plasmid loss probability as defined in (C). Statistics in (A) and (C) are for one-way ANOVA tests with the (A). plasmid transfer probability or (B) plasmid loss probability as the sole explanatory variable (n = 10).

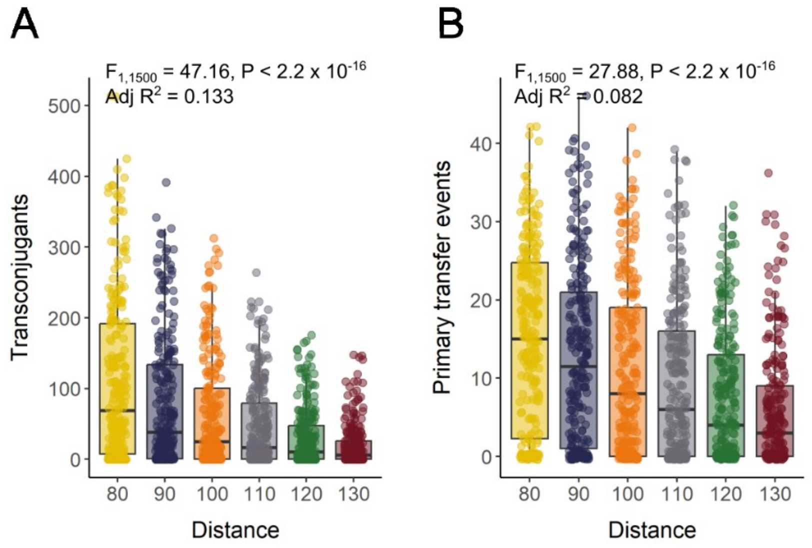


**Supplementary Figure 6.** Effect of the initial distance between colonies on plasmid spread at a fixed simulation time step of 640. (A) Total number of transconjugant cells (blue) as a function of the distance between inocula. (B) Total number of primary plasmid transfer events between the plasmid donor and potential recipient colonies as a function of the distance between inocula. Statistics are for one-way ANOVA tests with distance as the sole explanatory variable (n = 10).

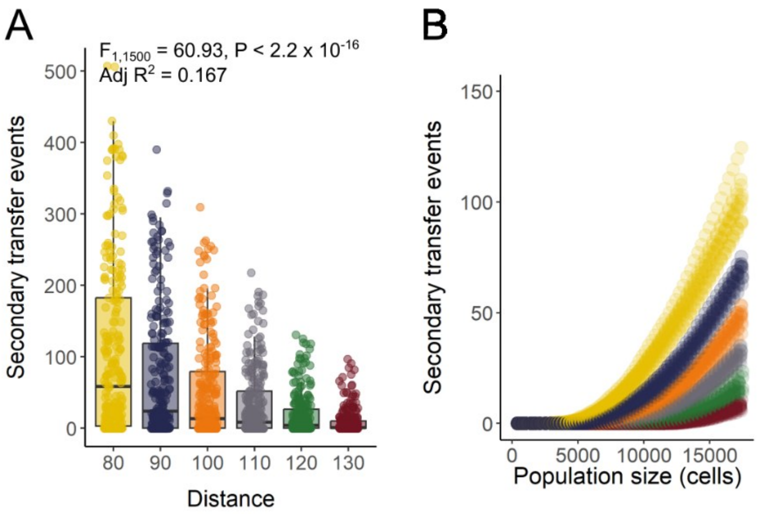


**Supplementary Figure 7.** Effect of the initial distance between colliding colonies on the total number of secondary plasmid transfer events within the potential recipient colony. (A) Total number of secondary plasmid transfer events at the end of the simulation as a function of the distance between inocula. (B) Total number of secondary plasmid transfer events as a function of the total population size. For (A), statistics are for one-way ANOVA tests with initial distance as the sole explanatory variable (n = 10).

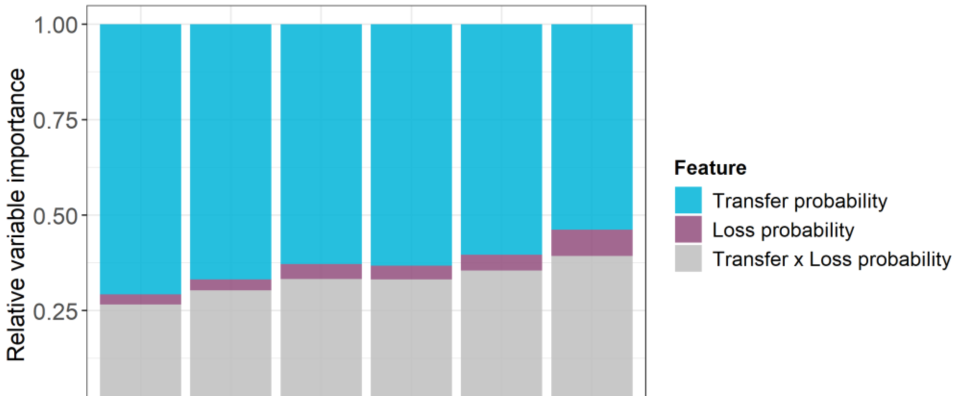


**Supplementary Figure 8.** Relative importance of the plasmid transfer and loss probabilities as a function of the initial distance between inocula on the total number of secondary plasmid transfer events within the potential recipient colony (n = 10).
